# Verticall: A fast and robust tool for recombination detection in large-scale bacterial genomic datasets

**DOI:** 10.64898/2026.04.21.719734

**Authors:** Erkison Ewomazino Odih, Ryan R. Wick, Kathryn E. Holt

## Abstract

The inference and removal of horizontally acquired genomic regions is a crucial step in phylogenomics analyses for evolutionary studies. Existing tools perform well on clonal lineage-focused datasets on the scale of hundreds of genomes, but are limited in their ability to analyse larger or more diverse datasets. Here we present Verticall, a tool to identify recombinant regions in bacterial assemblies and generate recombination-free phylogenies, which scales to thousands of genomes from clonal to genus-level diversity. Verticall uses a non-parametric approach to assign genomic regions as horizontally or vertically related based on the distribution of pairwise genetic distances between genomes. Recombination-free phylogenetic trees may be inferred by either calculating a pairwise genetic distance matrix from vertical-only regions (distance-tree approach) or by pairwise comparisons of all genomes to a reference and then masking horizontally acquired regions in a pseudo-alignment to the reference (alignment-tree approach). We demonstrate Verticall’s performance using four publicly available whole-genome sequence datasets of varying sample sizes (range: 154 – 4,857 genomes) and evolutionary scales (ranging from within-lineage to genus-wide diversity). Across all four datasets, Verticall showed comparable or superior performance to the established tools Gubbins and ClonalFrameML in terms of computational efficiency, plausibility of inferred phylogenetic trees, and recovery of temporal signal for molecular dating. Our results show that Verticall is a useful tool to more efficiently and accurately detect recombination, particularly applied to datasets for which existing tools are limited, including large datasets with hundreds to thousands of genomes and those that span entire species or genera. Verticall is available free and open source at https://github.com/rrwick/Verticall.

**Impact Statement:** Many bacterial species can acquire genetic material from external sources and stably incorporate them into their own genomes through homologous recombination. During phylogenomic analyses to investigate outbreaks or for evolutionary studies, a core objective is often to reconstruct the evolutionary history of the studied organisms independent of these horizontally acquired genomic regions. This is particularly desirable when the aim is to construct dated phylogenies, as horizontally acquired variation can interfere with the molecular clock signal on which dating relies. Existing recombination detection programs perform well in certain contexts, but their algorithms are not suitable for datasets with very high diversity or thousands of genomes. We addressed this gap by developing the software package Verticall. We show this approach produces comparable results to existing software for smaller more clonal datasets, but also performs well on datasets that the existing packages cannot handle.

**Data Summary:** Verticall is available free and open source at https://github.com/rrwick/Verticall. We used published whole-genome sequence data deposited in public databases (Pathogenwatch [https://pathogen.watch/]; European Nucleotide Archive [https://www.ebi.ac.uk/ena/], Sequence Read Archive [https://www.ncbi.nlm.nih.gov/sra/]). Accession numbers for the raw whole-genome sequences are presented in Tables S2–S6. All data, code, and analysis commands used to generate the results and figures presented in this paper are available on figshare (DOI: 10.6084/m9.figshare.31930821) and GitHub (https://github.com/erkison/verticall_paper).

## Introduction

Phylogenomics is the field of reconstructing evolutionary histories using whole-genome sequence data [1]. The resulting phylogenies can then be used for a wide range of analyses, including taxonomic classification [2], molecular dating [3] and outbreak investigation [4].

Using the entire genome (not just a select few genes) allows for greater phylogenetic resolution, especially with closely related organisms that have a small number of genomic differences. There exist a wide range of algorithms to conduct phylogenetic reconstruction from whole-genome data, including distance-based [5], maximum-likelihood [6] and Bayesian methods [7].

When bacteria evolve with 100% vertical inheritance (i.e., all DNA is transmitted from parent cell to daughter cells), all parts of the genome share the same evolutionary history. This makes phylogenetic reconstruction relatively straightforward, and many phylogenetic algorithms perform well in this scenario. However, many bacterial species also engage in horizontal gene transfer (HGT), acquiring genetic material from distantly related strains or even different species [8–10]. Bacterial HGT may occur through several mechanisms, including transformation, transduction, and conjugation, and may result in the transfer of small to large fragments of DNA, or even independently replicating mobile genetic elements like plasmids. To varying extents in different bacterial species, transferred DNA segments can be stably incorporated into the recipient’s genome through homologous recombination, driving rapid diversification and evolution [11–13].

The incorporation of genetic material at similar loci in the recipient genome via recombination is a major driver of bacterial evolution and adaptation, particularly in naturally competent bacterial species. From a phylogenomics standpoint, this presents a challenge as different regions of the genome may represent distinct evolutionary histories due to these recombination events. Including these recombined genome segments during phylogeny construction can introduce incongruent phylogenetic signals, causing erroneous or misleading tree topologies and/or branch lengths [14]. This is particularly important in outbreak investigations and epidemiological studies, where the inference of clonal relationships is crucial to accurately track transmission events and clonal evolution. Recombination can also obscure temporal signals of evolution in genomic data and bias estimates of substitution rate and divergence time [3]. The accurate detection and removal or exclusion of recombined genomic regions is thus a critical step in phylogenomic analyses.

Different tools are available for the detection and removal of recombination regions from whole-genome alignments prior to phylogenomic analysis, with Gubbins and ClonalFrameML being two of the most widely used. ClonalFrameML [15] identifies recombination regions based on the relative densities of polymorphisms along the alignment, while Gubbins [16] uses a similar iterative, phylogeny-aware approach based on significantly elevated mutation densities to detect recombination events on each branch. Both methods perform well for identifying and studying recombination regions within hundreds to a few thousand genomes of closely related strains, but are limited and indeed not recommended for the handling of large-scale genome datasets with extensive diversity [15,16]. Here, we present Verticall, a novel tool to identify and mask recombinant regions in large-scale and highly diverse genome datasets.

### Implementation and design

Verticall analyses genome assemblies to identify and mask regions likely acquired through horizontal transfer, retaining only the regions most consistent with vertical inheritance for calculating pairwise distances (distance-tree workflow) or generating a masked whole-genome pseudoalignment (alignment-tree workflow). It works by considering each pair of genomes in turn (**Figure 1**). A pairwise alignment is constructed, and genetic distance (nucleotide divergence) is calculated for sliding windows across the alignment to generate a distribution of distances for the genome pair. This distribution can be bi- or multi-modal if recombination has occurred in either lineage since the two genomes diverged, with the largest peak of genetic distances assumed to correspond to segments of the genome related through vertical evolution (diverging through clock-like accumulation of substitution mutations), and the other peak/s to segments of the genome that diverge through horizontal gene transfer. The observed distance distribution is partitioned into ranges associated with the largest peak (vertical evolution) vs others. This partitioning is used to (i) calculate the mean genetic distance associated with the largest peak, representing an estimate of nucleotide divergence by vertical evolution; and (ii) to assign individual segments of the genome as related by vertical or horizontal evolution based on genetic distance. Recombination-filtered trees can then be inferred, either from a matrix of pairwise vertical evolutionary distances (distance-tree workflow), or by comparing all genomes pairwise to a common reference, to generate a pseudo-alignment with non-vertically-inherited regions masked (alignment-tree workflow). Details of each of these steps are presented below.

**Figure 1:**
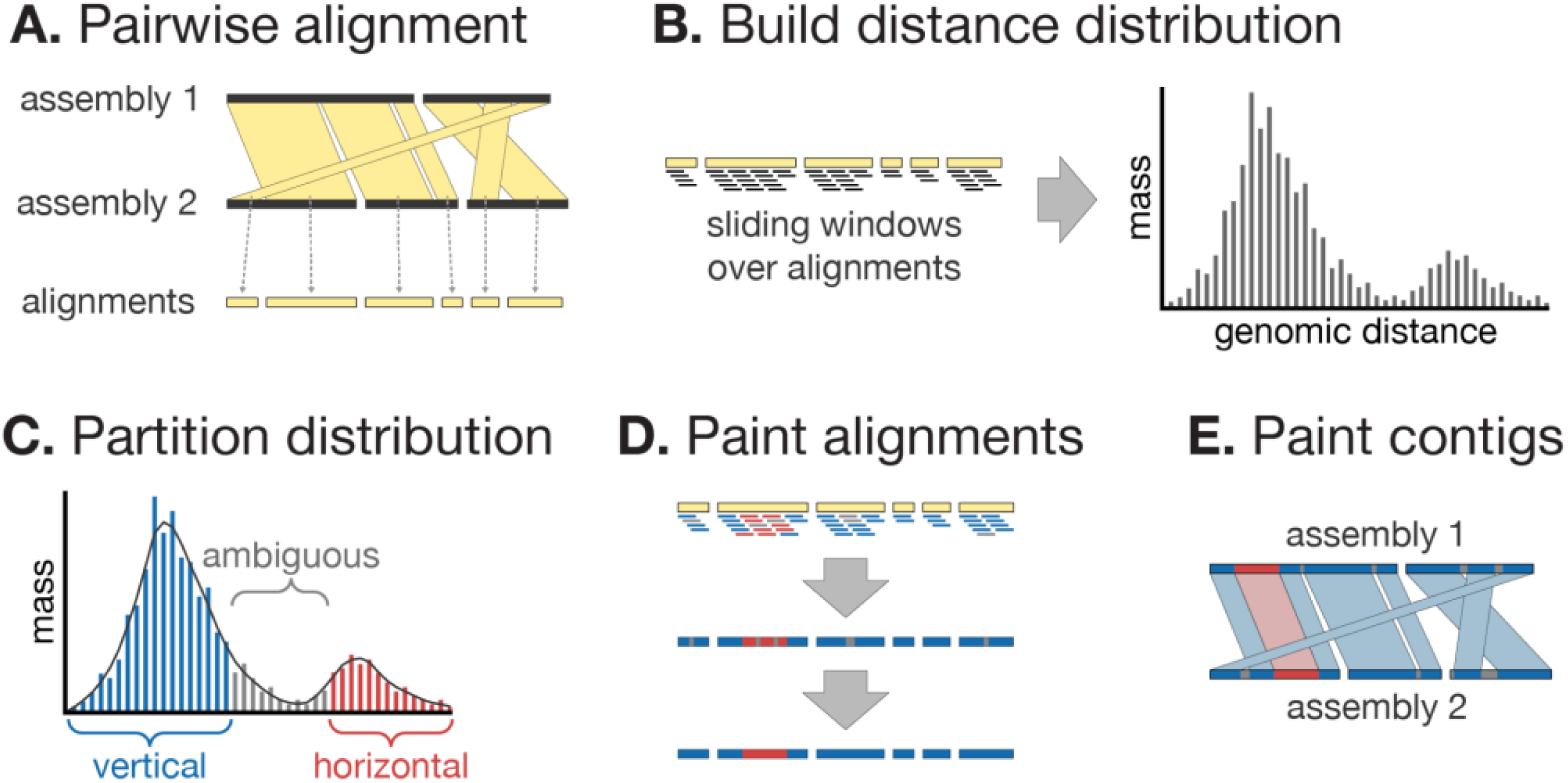
Verticall’s process for pairwise comparison of two assemblies. (A) The assemblies are aligned with minimap2. (B) A sliding window over the alignments is used to build a distribution of per-window difference counts. (C) This distribution is smoothed and partitioned, with the most massive peak assumed to represent vertically inherited sequence. (D) These classifications are painted onto the alignments and ambiguous regions are resolved according to their neighbours. (E) The painted alignments are mapped back onto the contigs, classifying regions as vertical, horizontal or unaligned.

### Pairwise assembly comparison

Pairwise assembly comparison, via the verticall pairwise command, is at the core of Verticall’s approach. For each pair of assemblies, Verticall first aligns them to each other using minimap2 [17] (**Figure 1A**). It then uses a sliding window over the resulting alignments to build a distance distribution (**Figure 1B**). This distribution represents how the genomic distance between the two assemblies varies depending on which part of the genome is compared, e.g. regions under purifying selection are expected to be on the low end of the distribution, while regions under diversifying selection are expected to be on the high end. Critically, if a region of the genome has a different evolutionary history to the rest (due to horizontal gene transfer), it may have an entirely different distance distribution, causing the overall distribution to be multimodal.

The user can provide an exact window size to use with the ‘--window_siz’ option, but if not given, Verticall will automatically choose a window size. It does this by aiming for a target number of windows (default = 50,000) and choosing a window size which allows for that many windows. This helps it to work under a wide range of conditions: when the alignments are short and highly fragmented (e.g. when the pair is distantly related), a shorter window is used, and when the alignments are long (e.g. when the pair is closely related and the assemblies are good), a longer window is used. The window step is 1% of the window size, e.g. if a window size of 10,000 bp is used, then adjacent windows will overlap by 9,900 bp.

Verticall then partitions the distribution into vertical, horizontal and ambiguous ranges (**Figure 1C**). It does this by first smoothing the distribution (using an Epanechnikov kernel [18]) and identifying all peaks in the smoothed distribution (ranges that extend from a local maximum to neighbouring local minima). The peak with the largest mass is considered to represent the vertical inheritance, and other peaks are considered to represent horizontal inheritance. The entire range of distribution is then partitioned as vertical, horizontal or ambiguous (ranges surrounding the minima between peaks).

Having partitioned the distance distribution, Verticall then uses these labels to ‘paint’ the alignments as vertical, horizontal or ambiguous (**Figure 1D**). Ambiguous regions are then resolved based on their neighbours: regions surrounded on both sides by vertical regions are changed to vertical, regions adjacent to a horizontal region are changed to horizontal, regions at the start/end of an alignment are changed to match their neighbouring region, and regions that span an entire alignment are conservatively changed to horizontal. This information can then be transferred back onto the two assemblies (**Figure 1E**), with each contig region ‘painted’ as vertical, horizontal or unaligned.

### Distance-tree workflow

The first way that Verticall can build a phylogeny is via the distance-tree workflow (**Figure 2A**). In this method, the verticall pairwise command performs pairwise assembly analysis on all possible assembly pairs (excluding self-vs-self pairs). This step scales O(n^2^) with the number of assemblies. Then the verticall matrix command can produce a PHYLIP distance matrix [19] of pairwise distances. Finally, an external program such as FastME [5] can be used to build a tree from the distance matrix.

**Figure 2:**
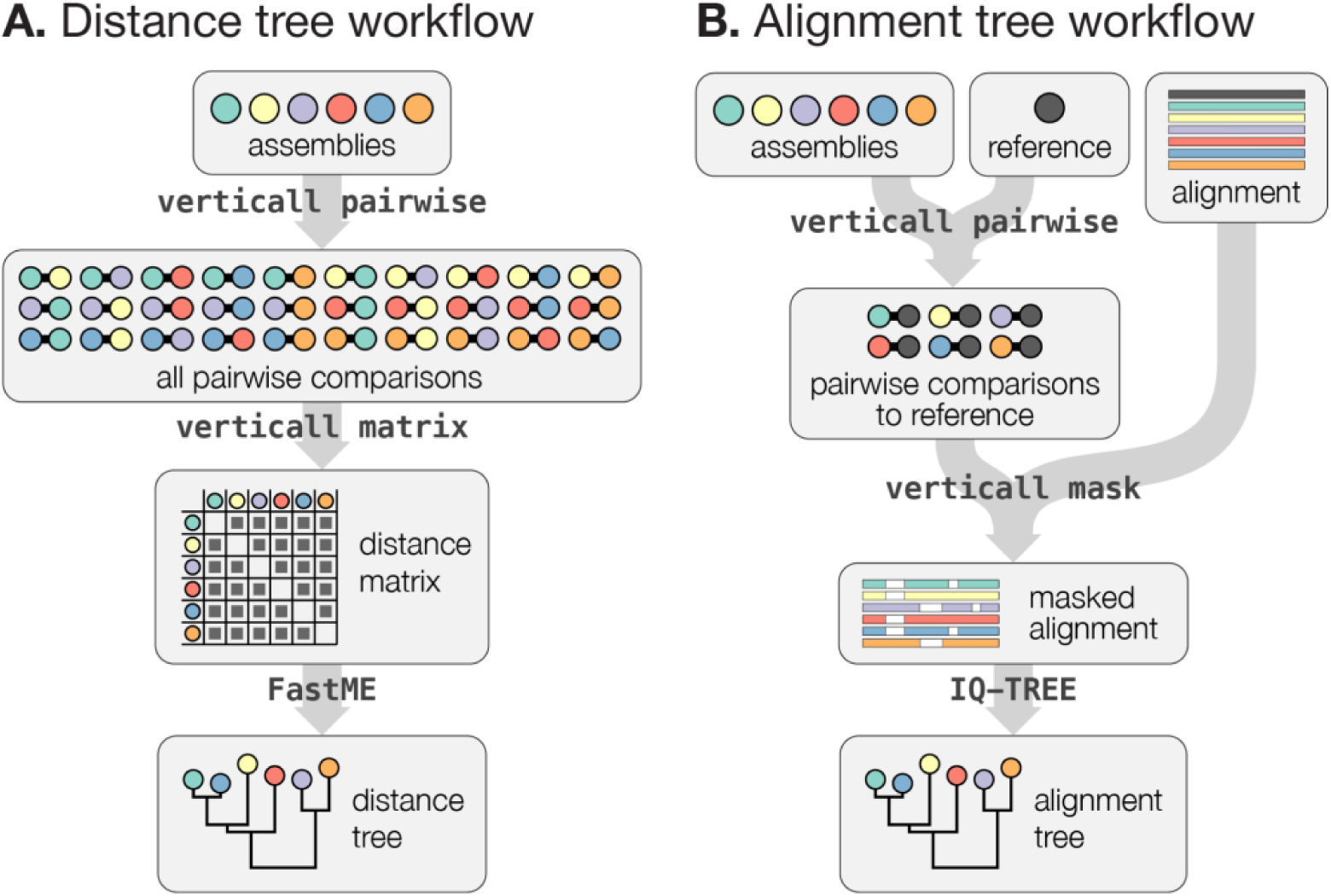
Overview of the two ways Verticall can be run: (A) a distance-tree workflow based on all pairwise comparisons between assemblies; and (B) an alignment-tree workflow based on comparing each assembly to a reference genome.

There are multiple ways one can determine a pairwise distance between two assemblies using the verticall pairwise results. The most useful is the interpolated median of the vertical component of the distance distribution. This metric serves to remove the effect of horizontal gene transfer in two ways. First, by only using the vertical component of the distance distribution, any regions painted as horizontal are not used. Second, the median is more robust to outliers than the mean, so even if verticall pairwise failed to identify a horizontally transmitted part of the genome, the median distance will be minimally affected. It is also crucial to use the interpolated median, not a true median, because when a pair of assemblies is very closely related, the true median distance is likely to be zero (**Figure S1**).

### Alignment-tree workflow

The second way Verticall can build a phylogeny is via the alignment-tree workflow (**Figure 2B**). In this approach, a whole-genome pseudoalignment based on a reference genome is required, and the verticall pairwise command performs pairwise assembly analysis between all assemblies and the reference genome. This step scales O(n) with the number of assemblies, making it much faster than the distance-tree workflow for large numbers of assemblies. Following this, the verticall mask command masks horizontal and unaligned regions of each genome in the pseudoalignment. Finally, an external program such as IQ-TREE [6] can be used to build a tree from the masked alignment. This approach is akin to the method used by other recombination-masking tools, such as Gubbins [16] and ClonalFrameML [15].

Compared to the distance-tree workflow, the alignment-tree workflow can be much faster due to its favourable computational complexity. It also allows for more robust tree-building algorithms, such as those based on maximum-likelihood or Bayesian algorithms [20]. However, the alignment-tree workflow only removes the effect of horizontal gene transfer in one way (masking horizontal and unaligned regions) while the distance-tree workflow additionally reduces its impact by using interpolated median distances.

### Code availability

Verticall is available free and open source under a GNU General Public License v3.0, at https://github.com/rrwick/Verticall. The version described in this paper is v0.4.2, DOI: 10.5281/zenodo.19687935.

## Methods

### Performance on datasets of different evolutionary scales

To test Verticall and other methods for building recombination-free trees, we used four publicly available datasets, each containing whole-genome Illumina reads (**Tables S1–S5**). The first contains 154 *Streptococcus pneumoniae* genomes from the PMEN1 lineage [21]. These genomes are closely related, have sample dates, contain a lot of horizontal gene transfer and have been previously used to test recombination detection methods [3,16]. The second dataset contains 4857 *Salmonella enterica* serovar Typhi genomes from the H58 lineage [22]. These genomes are closely related, have sample dates, contain a small amount of recombination and, due to the large number of genomes, make for a good test of computational resources. The third dataset contains 193 *Escherichia coli* genomes [23]. These species-wide genomes are diverse, with pairwise nucleotide divergences of up to ∼3.5%, and contain much recombination, including within-tree recombination. The fourth dataset contains 511 *Klebsiella* genomes spanning multiple species [24]. These genomes are very diverse, with pairwise nucleotide divergences of up to ∼16%, and contain much recombination, including cross-species hybridisation.

We assembled each genome using Unicycler v0.5.0 [25]. Genomes were then excluded based on the following criteria: lacking a sample year (for the *S. pneumoniae* and *S. enterica* datasets), N50 contig size of <15 kbp, assembly size of >7 Mbp, >100 assembly graph dead ends [26], or >1000 contigs. See **Tables S1–S5** for the details on the reads, assemblies and QC status of each genome.

For each dataset, we produced trees using both alignment-based and distance-based methods. For all alignment-based methods, we produced whole-genome pseudoalignments using Snippy v4.6.0 (https://github.com/tseemann/snippy) and built maximum-likelihood trees with IQ-TREE v2.3.6 [6]. We also attempted recombination filtering with Gubbins v3.4 [16], ClonalFrameML v1.13 [15] and Verticall’s alignment-tree workflow (version 0.4.1). We tested for the presence of recombination in the final alignments using the pairwise homoplasy index and maximum Chi-squared tests implemented in PhiPack v1.1 with 1000 permutations [27,28]. For distance-based methods, we produced a distance matrix with both FastANI [29] and Verticall’s distance-tree workflow, followed by tree-building with FastME [5]. All programs were run with default parameters except Gubbins for which the ‘--filter-percentag’ parameter was set to 100 to ensure that all samples were included.

Each program was given 32 CPU threads, 375 GB of RAM and one week (168 hours) of runtime. If the program did not complete (either due to RAM or time limits), we excluded that analysis for that dataset. The Gubbins and Verticall alignment-based trees for the *Klebsiella* dataset failed due to time. The *S. enterica* ClonalFrameML tree failed due to RAM, and the *S. enterica* Verticall distance-tree workflow failed due to time (see **Table S1**).

For all datasets, we assessed tree quality by quantifying root-to-tip variance. We used the FastRoot tool (v1.5) [30] to apply minimum-variance rooting to the tree and then quantified root-to-tip variance using the ape (v5.8) package in R. Lower root-to-tip variance was taken to indicate a more balanced tree [31]. For the *S. pneumoniae* and *S. enterica* (i.e. within-clone) datasets, we compared the recombination-filtering approaches by assessing their ability to recover temporal signal. We used the ape package (v5.8) [32] in R to best-fit root the tree, then performed a root-to-tip regression using the sample dates with the BactDating R package (v1.1.1) [3]. A stronger temporal signal (higher R^2^) was taken to indicate better recovery of molecular clock signal [8]. For the *E. coli* (species-wide) dataset, we also assessed each tree by comparing the pairwise patristic distances between and within clonal groups, and calculated the ratio of between-group to within-group distances (b/w ratio), where a higher ratio indicates a better phylogenetic delineation of clonal groups.

### Performance for dating analysis on >80 clonal lineages

We further evaluated the performance of Verticall and Gubbins by using each to analyse recombination and temporal signal within a fifth dataset of 83 *Klebsiella pneumoniae* lineages (**Tables S1, S6–S7**). We retrieved all 34,764 *K. pneumoniae* species complex genomes available on Pathogenwatch [33] as of June 2023, as well as 4,010 additional publicly available *K. pneumoniae* genomes from the KlebNET-GSP Genotype-Phenotype Group [34]. Non-*K. pneumoniae* genomes (n = 2,873) and genomes with suboptimal quality according to the KlebNET-GSP criteria (genomes with N50 < 20,000, contig count > 500, GC content < 56.35% or > 57.98%, total size < 4,969,898 bp or > 6,132,846 bp, or K or O locus confidence reported as ‘Low’ or ‘None’) (n = 5,247) were excluded from the analyses. Genomes were assigned to sublineages using core genome multi-locus sequence typing (cgMLST)-based Life Identification Number (LIN) codes [35]. This yielded 85 sublineages each with ≥20 genomes whose isolation dates spanned ≥10 years (n = 27,286 genomes total, see **Table S7**). A recently published collection [36] of high-quality, hybrid-assembled complete *K. pneumoniae* genomes sequenced using Illumina and Oxford Nanopore Technologies platforms were obtained for use as reference genomes for 34 of the selected sublineages (see **Table S8** for complete list of reference genomes used for each sublineage). For the remaining sublineages, we used Bactinspector v0.1.3 (https://gitlab.com/antunderwood/bactinspector) to select complete reference genomes from the RefSeq database [37]. No suitable reference genomes were identified in the RefSeq database for two of the sublineages (SL11518 and SL10226), so these were excluded from further analyses.

All genomes in each of the remaining 83 sublineages were mapped to their respective reference genomes using the Nextflow implementation of the RedDog pipeline (https://github.com/katholt/reddog-nf) with default parameters, and invariant sites were added to the final pseudoalignments before downstream analyses. The pseudoalignments were then used as input into Verticall alignment and Gubbins (run as previously described) for the detection and masking of putative horizontally acquired regions in the genomes. The Verticall distance workflow was also run directly on the assemblies as previously described. To reduce the runtime and computational resources required for the analyses, we downsampled sublineages with >200 genomes to reduce redundancy whilst maintaining the diversity within each sublineage. Downsampling was done using Treemmer (v0.3) [38] with constraint parameters added to keep at least 200 samples and at least one sample for every unique combination of K locus, O locus, and country and year of isolation.

We analysed the final trees output by the three recombination detection workflows, using the BactDating v1.1.1 [3] package in R (version 4.2.3) to assess temporal signal and estimate dated trees. The input into the *BactDate()* function was the final generated tree and the sampling years. Genomes missing sampling year information were initially included (years represented as ‘NA’) in the *BactDate()* runs but were excluded if the run failed to converge. The Gubbins tree output, along with all relevant recombination information, was imported using the *loadGubbins()* BactDating function. For each sublineage, we ran *BactDate()* for at least 10 million Markov Chain Monte Carlo (MCMC) iterations with sampling every N/1000 iterations after discarding the first half as burn-in. MCMC convergence was assessed based on trace plots of the sampled distributions. Runs with a stationary sampled distribution and with effective sample size (ESS) values >200 for all estimated parameters were deemed to have converged. We conducted all *BactDate()* runs using both the ‘relaxedgamma’ (relaxed clock) and ‘mixedgamma’ (mixture of the relaxed and strict molecular clock) models, and compared outputs from both models using the deviance information criterion (DIC). The model with the lower DIC was selected as the better fit for the data, except where any of the model estimates had ESS values <200, in which case we selected the model with the higher DIC and with ESS values ≥200 for all parameter estimates. We compared the mean substitution rate and root date estimates across all three recombination removal methods, by calculating the intra-class correlation (ICC) using a two-way mixed effects model with the irr v0.84.1 R package (https://cran.r-project.org/web/packages/irr/). We interpreted the ICC values as indicative of poor agreement (ICC < 0.5), moderate agreement (0.5 ≤ ICC < 0.75), good agreement (0.75 ≤ ICC < 0.9), or excellent agreement (ICC ≥ 0.9) between the compared methods [39]. Temporal signal and dated trees were also estimated for the *S. pneumoniae* dataset as described above. Based on the phylogenetic trees generated using all three methods, we excluded two genomes (SAMEA677487 and SAMEA677513) prior to the BactDating runs due to their high genetic dissimilarity (distinct branching and very long branch lengths) to the rest of the genomes.

## Results and Discussion

We explored Verticall’s performance by using it to generate recombination-filtered trees using the distance-based and alignment-based workflows, and comparing these to (i) recombination-filtered trees inferred using ClonalFrameML and Gubbins, and (ii) unfiltered trees inferred using distance-based or alignment-based approaches (see **Methods**). We attempted all six workflows to analyse four bacterial datasets, representing different population structures and evolutionary scales (within-clone *S. pneumoniae* and *Salmonella* Typhi, species-wide *E. coli*, and genus-wide *Klebsiella*), however some failed due to exceeding memory or runtime (see **Table 1**).

**Table 1:**
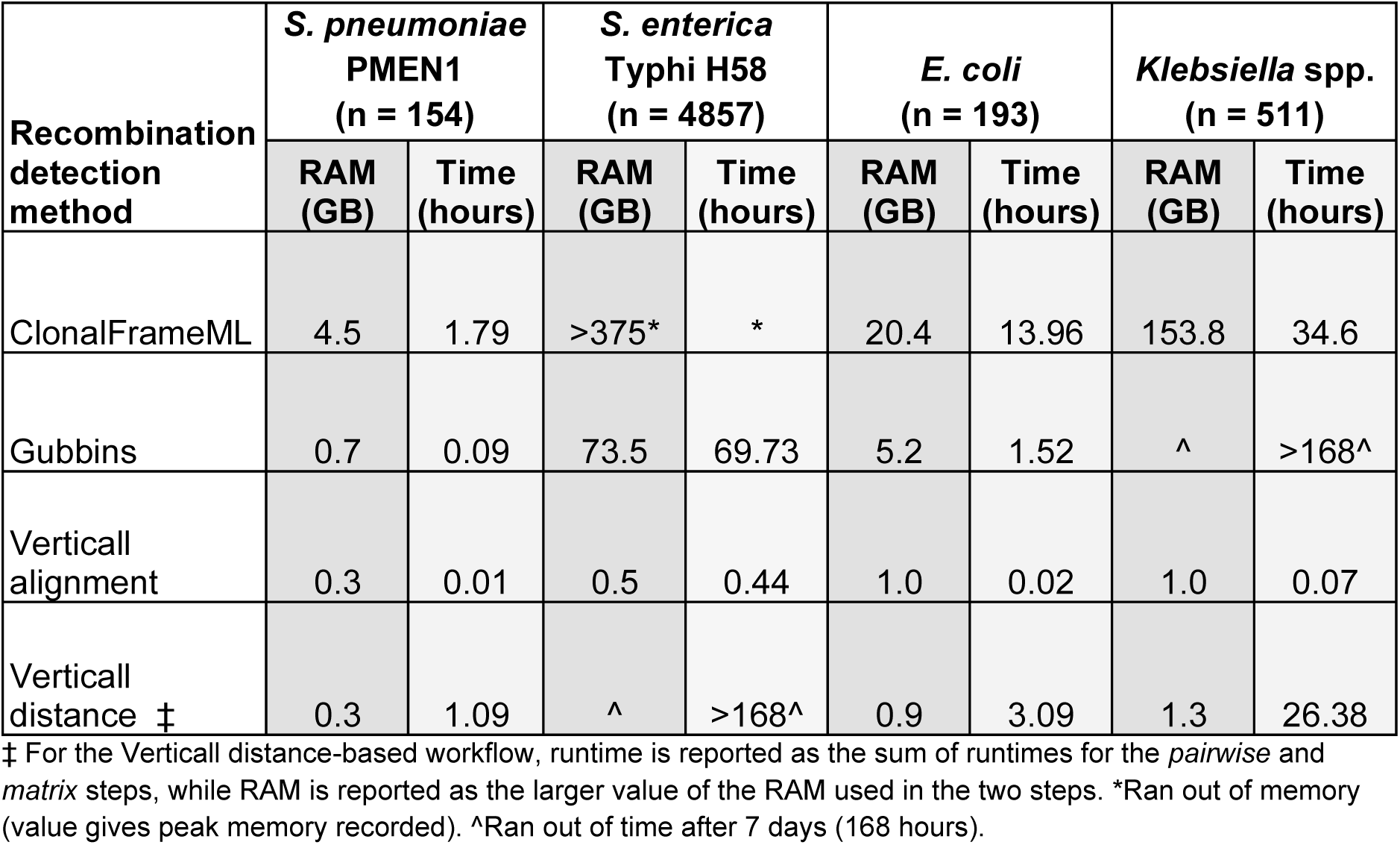
Computational resources used by each recombination detection method.

All six workflows completed for the S. *pneumoniae* PMEN1 clone dataset, which was used in the original assessment and validation of Gubbins [16]. There was significant evidence of recombination in the unfiltered alignment based on the spatial distribution of polymorphic sites (maximum Chi-squared test, *p* = 0) and the presence of homoplasies (pairwise homoplasy index [PHI], *p* = 0). The Verticall alignment workflow identified and masked a total of 560 recombination blocks across all 154 genomes (median: 2 blocks per genome, interquartile range [IQR]: 0–4, range: 0–24; block size range: 1,670 – 26,425 bp) (**Figure 3A**), but there was still significant evidence of recombination in the masked alignment (maximum Chi-squared test, *p* = 0 and PHI, *p* = 0). Using ClonalFrameML, 2,569 recombination events in total were identified (**Figure 3B**), with recombination block sizes ranging between 2 and 24,311 bp (median = 1,410 bp, IQR: 635–3,538 bp). As with the Verticall alignment workflow, the PHI (*p* = 0.031) and maximum Chi-squared (*p* = 0.023) tests also showed significant evidence of residual mosaicism in the ClonalFrameML-filtered alignment. Following recombination detection and removal using Gubbins, which predicted 2,601 recombination events in total (median: 17, IQR: 11–20, range 0–41) (**Figure 3C**), there was no evidence of recombination either based on the PHI test (*p* = 0.27) or maximum Chi-squared test (*p* = 0.058). The predicted recombination regions ranged in size between 3 and 43,563 bp (median = 3,538 bp, IQR: 722–7,881 bp). Verticall distance (distance-tree workflow) identified similar regions of recombination compared to Gubbins (**Figure 3D**), but also predicted thousands more putative recombination regions not identified by Gubbins (n=15,980 [median: 107, IQR: 98–114, range: 32–127]; block size range: 76–73,927 bp), which is expected given the pairwise architecture of this method (overlapping recombination blocks detected relative to ≥2 other genomes were merged and counted once to reduce redundancy). The statistical tests for recombination could not be conducted for the Verticall distance workflow as it does not produce an alignment. Gubbins identified four distinct clades (coloured in **Figure 3**) delineated by major recombination blocks; the Verticall alignment tree resolved three of these clades, while ClonalFrameML and the Verticall distance trees resolved all four. Nevertheless, there were also some notable differences between the overall topologies of the Gubbins and Verticall distance trees, including trivial differences resulting in varied placements of certain genomes within distinct clades, as well as a few marked differences disrupting predicted clonal relationships. For instance, Gubbins grouped isolates SAMEA677529, SAMEA677516, SAMEA677524, and SAMEA677405 along with 12 other isolates in a single clade (all coloured red in **Figure S2**). These four isolates were however separated by the Verticall distance workflow into a more ancestral node on the tree. Verticall required less RAM (0.3 GB for each workflow) compared to Gubbins (0.7 GB) and ClonalFrameML (4.5 GB), and finished in a shorter time compared to the others (Verticall alignment = 0.01 hours, Gubbins = 0.09 hours, Verticall distance = 1.09 hours, ClonalFrameML = 1.79 hours) (**Table 1**).

**Figure 3:**
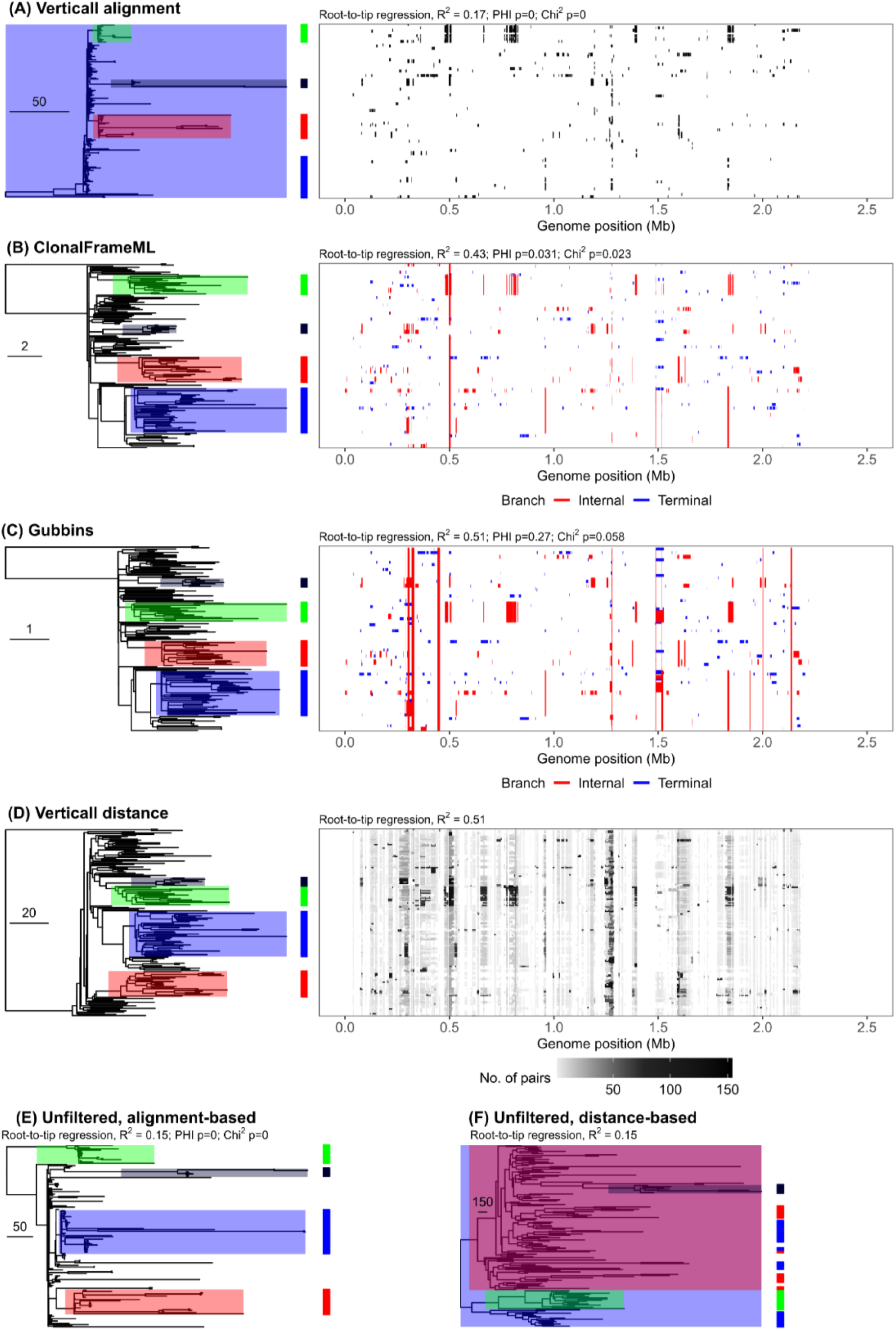
Comparison of the Gubbins, ClonalFrameML, and Verticall workflows for the analysis of *S. pneumoniae* PMEN1 lineage sequences. Plots show the recombination blocks (right subpanels) and final trees (left subpanels) produced using the Verticall alignment **(A)**, ClonalFrameML **(B)**, Gubbins **(C)**, and Verticall distance **(D)** workflows. Unfiltered trees generated using alignment-based **(E)** and distance-based **(F)** approaches are also shown. The tree branches are shaded to indicate four clades delineated by major recombination blocks in the Gubbins tree and the nodes representing the most recent common ancestors of the corresponding genomes in the Verticall alignment, ClonalFrameML, Verticall distance, and unfiltered trees. Individual genomes belonging to each of the four clades are indicated in the adjoining strip next to each tree. Tree scales indicate number of substitutions. Each row in the recombination plots (right subpanels) represents a single genome, and the columns correspond to nucleotide positions in the reference genome (*S. pneumoniae* ATCC 700669 chromosome; GCF_000026665.1). In panels (B) and (C), the recombination blocks are coloured to indicate recombination regions predicted in internal (shared by multiple genomes) or terminal branches, as per the panel legends. In (D), the blocks shown are pairwise recombination blocks predicted by the Verticall distance workflow relative to other genomes in the dataset. Overlapping recombination blocks were merged and the blocks are coloured to indicate the number of genome pairs relative to which the recombination regions were detected. Reference-based positions for each block were determined by aligning each genome to the reference; blocks outside reference-aligned positions are not shown (n=530/15,980). Tests for evidence of recombination (PHI and Chi^2^) are shown for the alignment-based workflows. PHI – pairwise homoplasy index (tests for presence of homoplasies); Chi^2^ – maximum Chi-squared test (spatial distribution of polymorphic sites).

For the *S. enterica* Typhi H58 dataset, which represents a large, highly clonal dataset (n = 4,857 genomes), ClonalFrameML and the Verticall distance workflow failed to complete due to memory and time, respectively. **Figure 4** shows the recombination regions detected by Gubbins and Verticall alignment in the *S. enterica* Typhi H58 genomes (runs to test for evidence of recombination in the unfiltered and recombination-filtered alignments failed to complete after 7 days). Both workflows detected a median 3 recombination regions across all genomes (Gubbins IQR: 2–4, range: 0–9 and Verticall alignment IQR: 3–3, range: 0–7). Verticall alignment predicted three major recombination blocks in at least 3,677 of the 4,857 genomes, one of which overlapped with a recombination block (∼6 kb) identified by Gubbins in 4,612 genomes. Both methods produced trees with similar topologies, with minor differences in the placement of closely related isolates within distinct clades (see **Figure 4**). The Verticall alignment workflow had a considerably quicker runtime (0.44 vs 69.7 hours) and required substantially less RAM (0.5 vs 73.5 GB) compared to Gubbins.

**Figure 4:**
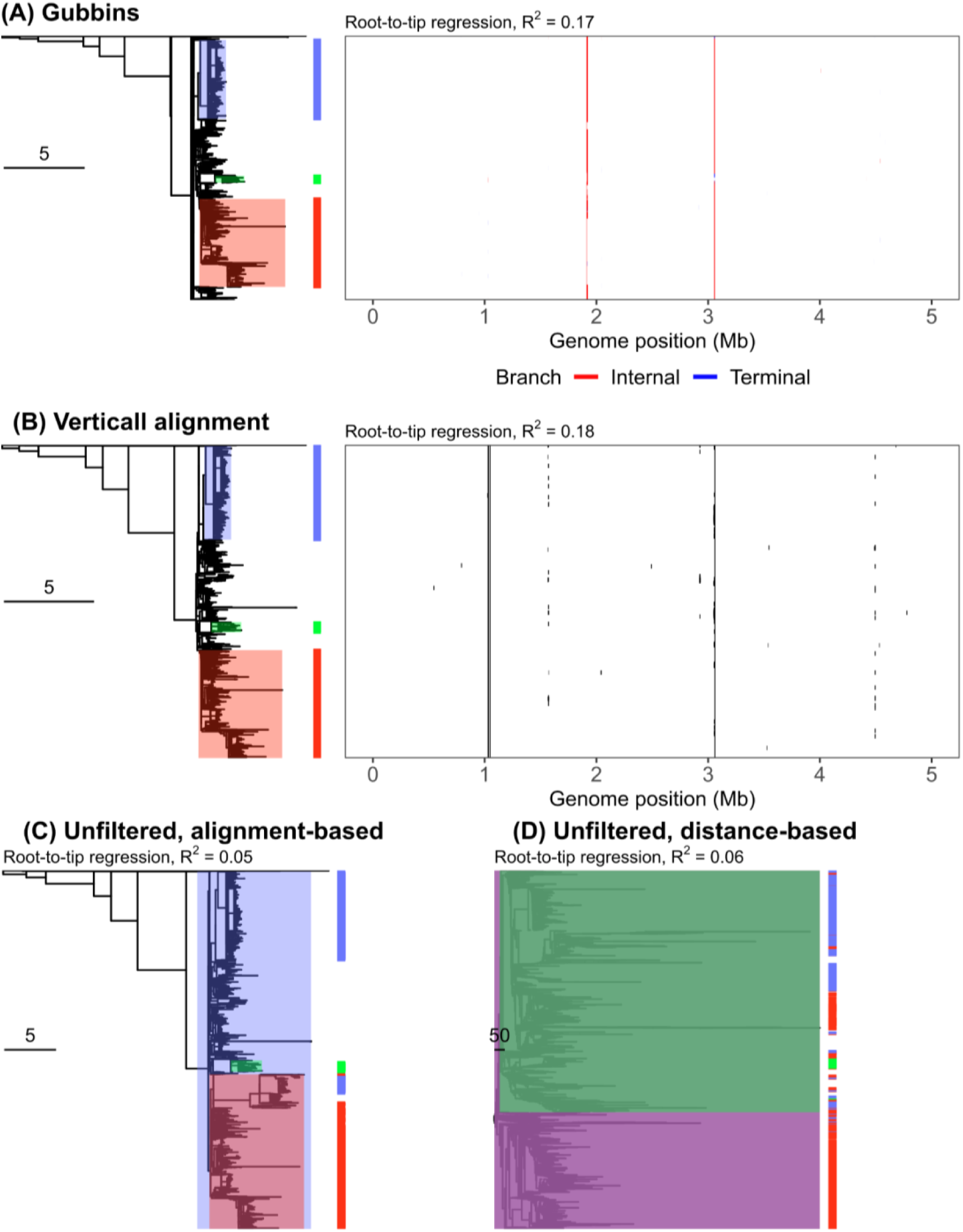
Comparison of the Gubbins and Verticall alignment workflows for the analysis of *S. enterica* Typhi H58 lineage sequences. Plots show the recombination blocks (right subpanels) and final trees (left panels) produced using the Gubbins **(A)** and Verticall alignment **(B)** workflows. Unfiltered trees generated using alignment-based **(C)** and distance-based **(D)** approaches are also shown. The tree branches are shaded to indicate three major clades in the Gubbins tree and the nodes representing the most recent common ancestors of the corresponding genomes in the Verticall alignment and unfiltered trees. Individual genomes belonging to each of the three clades are indicated in the adjoining strip next to each tree. Tree scales indicate number of substitutions. Each row in the recombination plots (right subpanels) represents a single genome, and the columns correspond to nucleotide positions in the reference genome (*S. enterica* Typhi CT18 chromosome; GCF_000195995.1). In panel (A), the recombination blocks are coloured to indicate recombination regions predicted in internal (shared by multiple genomes) or terminal branches, as per the panel legends.

Next we assessed performance on the *E. coli* species-wide dataset (n = 193 genomes), for which all six workflows completed successfully. After recombination filtering, there was still significant evidence of recombination in the alignments produced by the Verticall alignment workflow (maximum Chi-squared test, *p* = 0 and PHI, *p* = 0), Gubbins (maximum Chi-squared test, *p* = 0 and PHI, *p* = 0), and ClonalFrameML (maximum Chi-squared test, *p* = 0.04 and PHI, *p* = 0); the test with the unfiltered alignment timed out after 168 hours. As ClonalFrameML was designed to analyse both species-wide and within-clone evolution [15], whilst Gubbins was designed only for within-clone analyses, we compare results primarily to ClonalFrameML as the reference method. The unfiltered trees showed high root-to-tip variances (distance-based: 0.0476, alignment-based: 0.052), reflecting imbalanced branch lengths influenced by different amounts of recombination in different parts of the tree. The ClonalFrameML tree was more balanced (variance = 0.0374), as were those produced by Gubbins (0.0236) and Verticall distance (0.0349). The Verticall alignment tree had a variance closer to the unfiltered trees (0.0443), reflecting poor performance in detecting and masking recombinant regions (see **Figure 5**). Consistent with this, the Verticall alignment workflow identified only 3 recombination blocks on average across all genomes (median: 3, IQR: 0–11, block size median = 7838 bp [IQR: 3927–13096 bp]), compared to a median 186 recombination events predicted by Verticall distance (median: 186, IQR: 160–213, block size median = 9438 bp [IQR: 5110–17427 bp]). Gubbins (median: 1616, IQR: 1388–1678, block size median = 390 bp [IQR: 141–1017 bp]) and ClonalFrameML (median: 2665, IQR: 2550–2962, block size median = 504 bp [IQR: 219–1123 bp]) each predicted over 1000 more recombination regions, albeit with comparatively smaller block sizes. The branching structure of the Verticall distance and ClonalFrameML trees were almost identical, with limited crossover of closely related isolates within distinct clades (see **Figure S3**). Next we assessed each tree in terms of its differentiation of distinct clonal groups (coloured in **Figure 6**), based on between- vs within-group patristic distances. The unfiltered alignment-based and distance-based trees showed poor differentiation (b/w ratios of 28.3 and 7.2, respectively), whereas ClonalFrameML- and Gubbins-filtered trees showed better differentiation (b/w ratios 37 and 67.8, respectively) and Verticall methods showed equivalent improvement (48.7 and 82.1) (see **Figure 6**). Verticall required considerably less RAM (≤1 GB for both workflows) and a shorter runtime (alignment workflow: 0.02 hrs, distance-tree workflow: 3.1 hours) compared to Gubbins (5.2 GB RAM, 1.5 hours) and ClonalFrameML (20.4 GB RAM, 14.0 hours) (**Table 1**).

**Figure 5:**
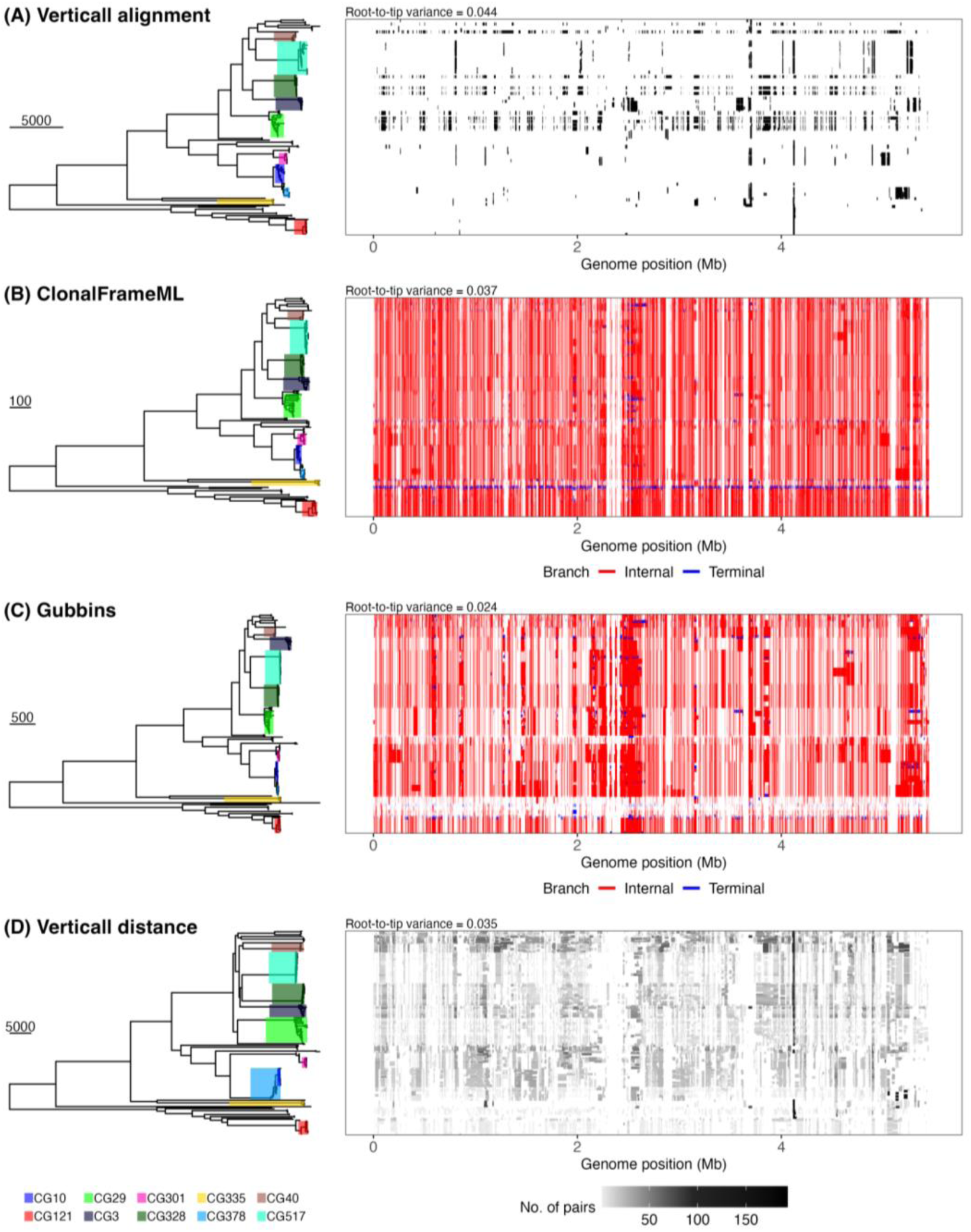
Comparison of the Gubbins, ClonalFrameML, and Verticall workflows for the analysis of *Escherichia coli* sequences. Plots show the recombination blocks (right subpanels) and final trees (left subpanels) produced using the Verticall alignment **(A)**, ClonalFrameML **(B)**, Gubbins **(C)**, and Verticall distance **(D)** workflows. The tree branches are shaded to indicate the different *E. coli* clonal groups, as per the figure legend. Tree scales indicate number of substitutions. Each row in the recombination plots (right subpanels) represents a single genome, and the columns correspond to nucleotide positions in the reference genome (*E. coli* O103:H2 12009 chromosome; GCF_000010745.1). In panels (B) and (C), the recombination blocks are coloured to indicate recombination regions predicted in internal (shared by multiple genomes) or terminal branches, as per the panel legends. In (D), the blocks shown are pairwise recombination blocks predicted by the Verticall distance workflow relative to other genomes in the dataset. Overlapping recombination blocks were merged and the blocks are coloured to indicate the number of genome pairs relative to which the recombination regions were detected. Reference-based positions for each block were determined by aligning each genome to the reference; blocks outside reference-aligned positions (n=8,495 / 34,985) are not shown.

**Figure 6:**
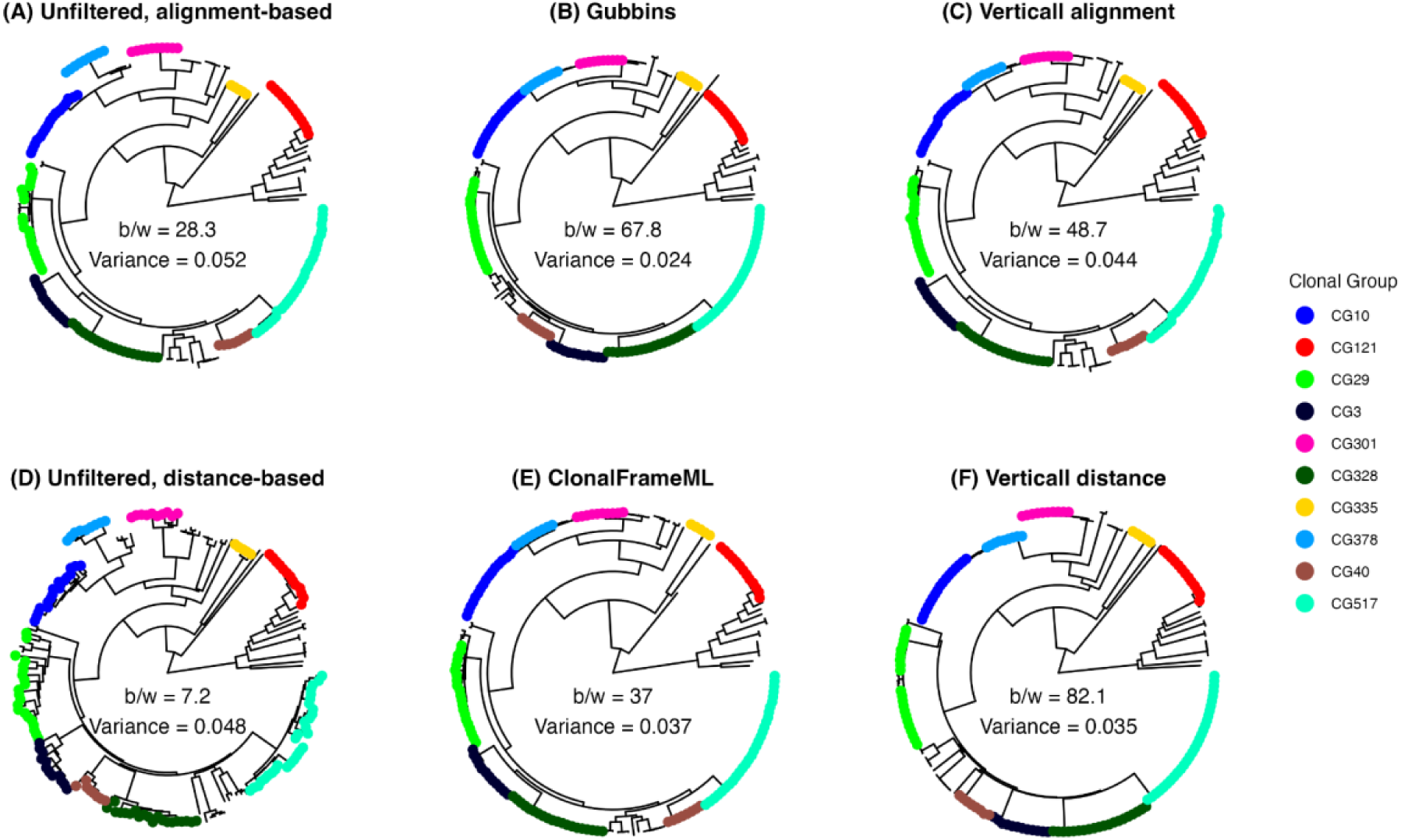
*Escherichia coli* phylogeny produced using different phylogeny construction workflows. Tree tips are coloured by the clonal group to which the strains belong, and the ‘b/w’ values show the ratio of between-clonal group to within-clonal group mean pairwise patristic distances. Root-to-tip variance values are also shown.

Finally, we applied all six workflows to a moderately large and highly diverse, genus-wide set of *Klebsiella* genomes, a type of dataset for which neither Gubbins nor ClonalFrameML are well suited or recommended. Only the unfiltered trees, ClonalFrameML, and Verticall distance workflows completed, with the others timing out after one week (Gubbins and Verticall alignment). Expectedly, ClonalFrameML showed poor performance (over-filtering) with this dataset, with over 2 million recombination regions predicted across all 511 genomes (median = 3,961 [IQR: 3,838–4,166]) and only 159 core genome sites left in the filtered alignment. Consequently, the inferred phylogenetic tree contained multiple polytomies and implausible branch lengths and failed to resolve the distinct species (**Figure S4**). Verticall distance predicted 40,538 total recombination regions (median = 80 [IQR: 69–91]), with recombination block sizes ranging from 16–1,352,787 bp. The Verticall distance tree showed the least variance in root-to-tip distances (0.014), compared with ClonalFrameML (0.116) and the unfiltered distance-(0.023) and alignment-based workflows (0.331) (**Figure S4**).

### Divergence time estimation using Verticall

A key use of recombination-filtered trees is to restore temporal signal to allow dating analysis, by identifying and excluding regions related through horizontal transfer to leave only those related through vertical evolution according to a measurable molecular clock rate [40]. We explored how well we can recover temporal signal within clones using Verticall in different modes, and compared to Gubbins. For the *S. pneumoniae* PMEN1 and *S. enterica* Typhi H58 clone datasets, we first assessed temporal signal based on correlation between root-to-tip distances and sampling dates. Our results show that Verticall distance produces similar results to Gubbins and superior results to ClonalFrameML in restoring temporal signal in a small dataset of closely related *S. pneumoniae* PMEN1 genomes, the kind of dataset for which Gubbins is especially well suited. For this dataset, the two unfiltered trees showed weak temporal signal (alignment-based: R^2^ = 0.15; distance-based: R^2^ = 0.15; **Figure 3, Figure S5**). The recombination-filtered trees produced by Gubbins and ClonalFrameML showed much stronger temporal signal (R^2^ = 0.51 and 0.43, respectively), consistent with the desired goal of masking out regions related through horizontal transfer to leave those related through vertical evolution according to a measurable molecular clock.

The tree produced by the Verticall distance workflow showed an equivalent clock-like signal to the Gubbins tree (R^2^ = 0.51), however the Verticall-alignment workflow failed to recover any greater temporal signal than was evident in the unfiltered trees (R^2^ = 0.17). For the *S. enterica* Typhi H58 dataset, the unfiltered trees showed no temporal signal (alignment-based: R^2^ = 0.05; distance-based: R^2^ = 0.06; **Figure 4, Figure S6**). The recombination-filtered trees generated by Gubbins and the Verticall-alignment tree workflow recovered moderate temporal signal (R^2^ = 0.17 and 0.18, respectively; see **Figure 4**), with Verticall requiring significantly less memory and a shorter runtime (0.5 vs 74 GB RAM; 0.4 vs 70 hours). ClonalFrameML and Verticall distance failed to complete due to running out of memory or time, respectively.

Didelot and colleagues [3] previously analysed the *S. pneumoniae* PMEN1 dataset described above using Gubbins for recombination correction, followed by analysis of the resulting recombination-filtered tree using BactDating for divergence date estimation. They estimated a substitution rate of 3.09 [95% credible interval (CI): 2.68–3.53] substitutions per genome per year, and a root date of 1972 [95% CI: 1966–1977]. Our analysis here used a subset of the same genomes, yielding root date estimates that were in excellent agreement with those previously published (1974 [95% CI: 1970–1977]), and a slightly lower substitution rate estimate (2.63 substitutions/year [95% CI: 2.25–2.99], see **Figure 7A**). Using the Verticall distance tree as input to BactDating analysis, our root date estimate (1970 [95% CI: 1964–1975]) was also in excellent agreement with Gubbins, but the mean substitution rate estimate obtained was lower (1.96 substitutions/year [95% CI: 1.65–2.30]), suggesting that Verticall may be underestimating the evolutionary distance (over-filtering; median recombination events per genome = 107 vs 17 with Gubbins; see **Figure 3D**). With the Verticall alignment workflow however, the estimated substitution rate was substantially lower (1.05 substitutions/year [95% CI: 0.71–1.49]) and the root date much earlier (1701 [ 95% CI: 1186–1915]), consistent with insufficient filtering (median recombination events = 2 vs 17 with Gubbins; see **Figure 3A**). We attempted to analyse the *S*. Typhi H58 Gubbins and Verticall alignment trees using BactDating, but the runs failed to converge after 10^6^ iterations, and attempting to run for 10^7^ timed out after 7 days. These examples suggest that Gubbins is best-suited to small sets of genomes with substantial amounts of recombination, whereas Verticall offers substantial advantages in computational efficiency and speed for large datasets with closely related genomes, while maintaining accuracy (identical R^2^ values for the 4,857-genome *S. enterica* Typhi dataset, in <1 hour compared with ∼70 hours; see **Figure S6**, **Table 1**).

**Figure 7:**
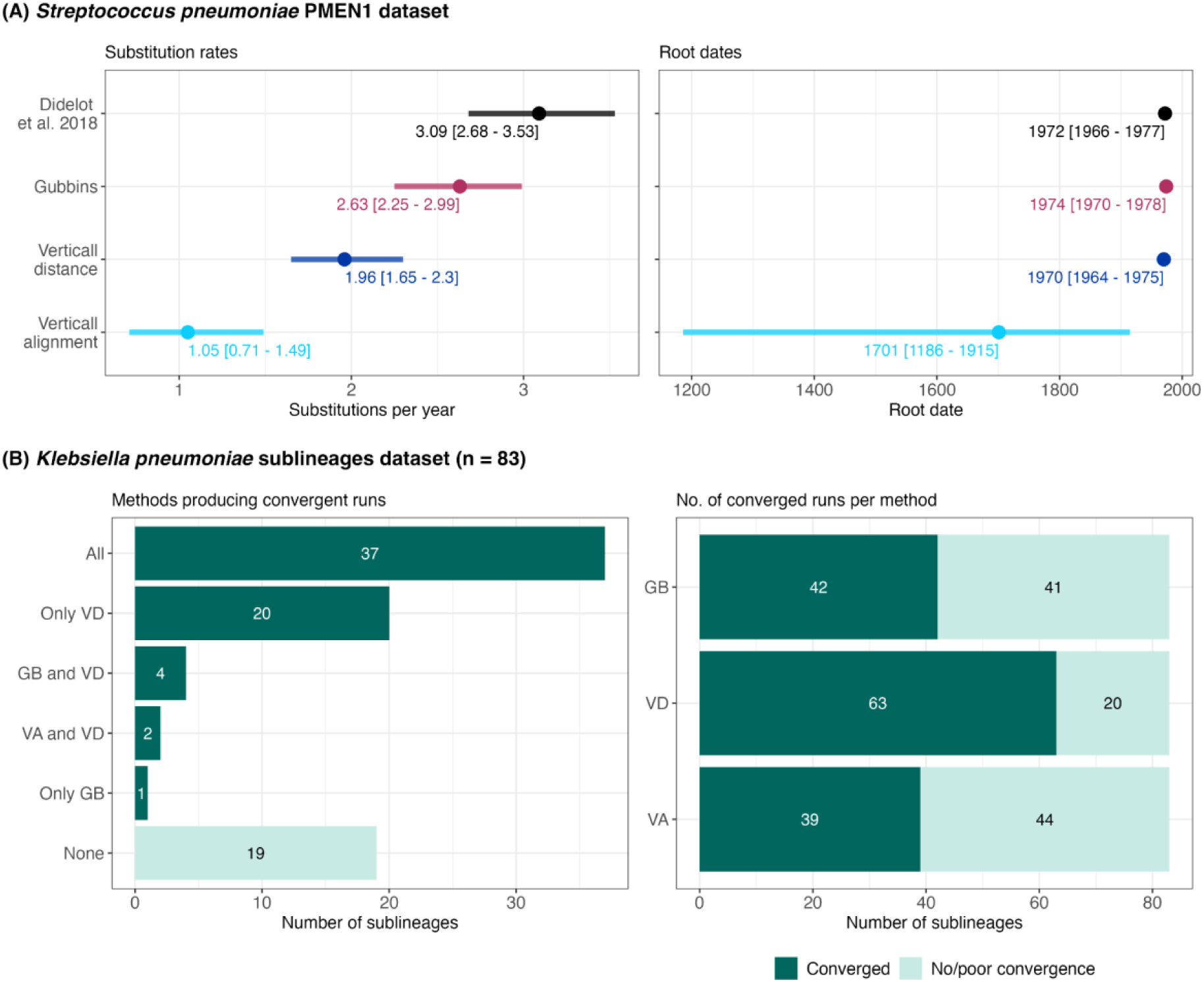
Comparison of Verticall and Gubbins workflows for divergence time estimation analysis. (A) *S. pneumoniae* PMEN1 lineage substitution rate and root date estimates obtained with BactDating using Gubbins and Verticall input trees (B) Performance of Verticall and Gubbins for the restoration of temporal signal in 83 *K. pneumoniae* sublineages for divergence time estimation analysis. GB - Gubbins, VA - Verticall alignment, VD - Verticall distance

To explore in more detail the reliability and robustness of Gubbins and Verticall as methods to filter recombination from within clones to restore temporal signal for dating analysis, we applied the three workflows to analyse 83 different *K. pneumoniae* clonal sublineages, with sample sizes ranging from 19 to 838 genomes (see **Methods**). Recombination-filtered trees were successfully generated by all three workflows for 83 lineages, and analysed using BactDating (**Tables S9–S11**). For 64 sublineages, BactDating MCMC runs converged using at least one of the three input trees (**Figure 7B**). For 37 sublineages, runs converged using all three trees. For a further 26 sublineages, dating succeeded using the Verticall distance tree, either alone (n=20) or together with Verticall alignment (n=2) or Gubbins (n=4); and for one sublineage (SL502) only succeeded using the Gubbins tree. MCMC runs for nineteen clones failed to converge using any of the three input trees. The Verticall distance workflow had the best performance of the three methods, recovering temporal signal in 63 (75.9%) of the 83 clones (**Figure 7B**). The Verticall alignment workflow (n=39/83; 47.0%) and Gubbins (n=42/83; 50.6%) had comparably lower performance, with each recovering temporal signal in around half of all *K. pneumoniae* sublineages analysed (**Figure 7B**).

To explore potential predictors of convergence (binary outcome), we fit multivariable logistic regression models for each of the three workflows where predictors were encoded as numerical variables (number of genomes, nucleotide divergence, root-to-tip regression coefficient [R^2^]). Although datasets with lower nucleotide divergence converged more frequently across all methods (see **Figure S7**), nucleotide divergence was not significant in any of the multivariable models, nor was root-to-tip R^2^ (**Table S12**). For the Verticall distance method, which had the highest rate of convergence, larger sample size was beneficial such that datasets with more genomes had significantly higher odds of convergence compared to smaller datasets (OR = 10 [1.71–58.31], *p* = 0.0105).

Among the 37 clones for which all three methods produced convergent runs, we excluded three clones for which at least one workflow produced negative root date estimates (SL34 – all three workflows, SL35 – all three workflows, and SL111 – Gubbins and Verticall alignment) and compared the estimated parameters (substitution rates and root dates) in the remaining 34 clones. Compared with Gubbins trees, Verticall distance trees generated overlapping substitution rate and root date estimates for all 34 clones (100%) (see **Figure 8, Tables S9–S11**). Overall agreement between estimates based on the Gubbins and Verticall distance trees was strong (ICC = 0.87 [95% CI: 0.75–0.93]) for substitution rates, but more moderate for root dates (0.51 [0.23–0.72]). Agreement between Gubbins and Verticall alignment trees was relatively poorer for substitution rates (0.45 [0.15–0.68]) and stronger for root dates (0.73 [0.53–0.86]).

**Figure 8:**
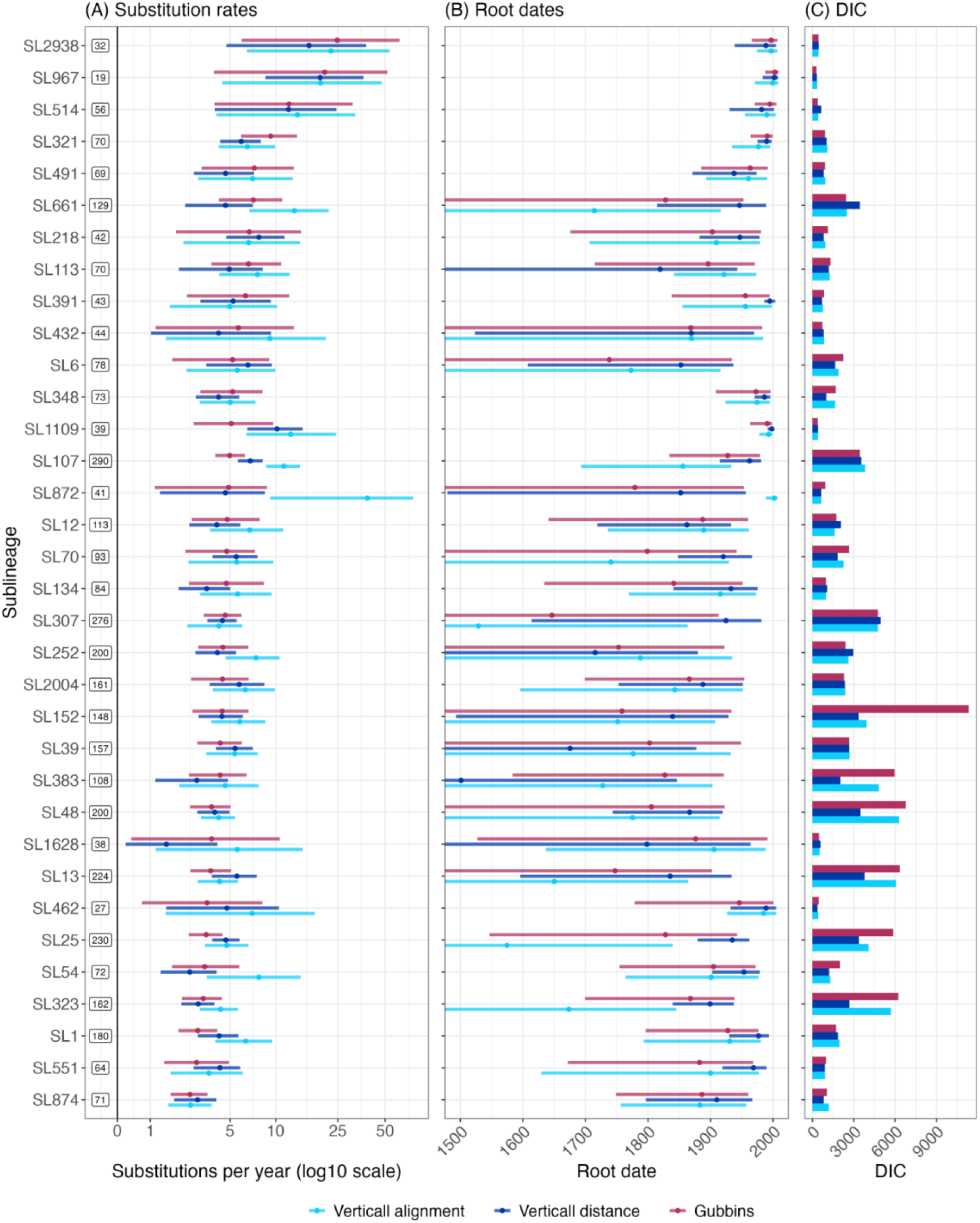
Comparison of molecular dating estimates obtained using Verticall- and Gubbins-filtered trees of 83 *K. pneumoniae* sublineages. (A) Substitution rate estimates (points) and 95% credible intervals (lines) estimated using the Gubbins, Verticall distance, and Verticall alignment recombination detection workflows, coloured as per the figure legend. Boxed labels indicate the number of genomes per sublineage. (B) Root date estimates. Lower bounds of the 95% credible interval exceeding year 1500 are truncated in the plot. See **Tables S9–S11** for all estimates and 95% credible intervals. (C) Deviance information criterion values for each model.

Published estimates of *K. pneumoniae* mutation rates, based on BEAST2 analysis of smaller datasets (N <100 each), were in the range 1.9 to 8.2 substitutions/year [95% CI: 1.3–10.0] [41–45]. Most mutation rates estimated here using BactDating analysis were in this confidence interval range, including 90.5% (n=38/42) of those estimated from Gubbins trees, 93.8% (n=60/64) of those from Verticall distance trees, and 82.1% (n=32/39) of those from Verticall alignment trees. Some notable clones with mutation rates outside this range include SL514 (Gubbins = 12.2 substitutions/year; Verticall distance = 12.1; Verticall alignment = 13.8), SL967 (Gubbins = 20.6 substitutions/year; Verticall distance = 19.3; Verticall alignment = 19.4), and SL2938 (Gubbins: 24.8 substitutions/year; Verticall distance: 16.4; Verticall alignment: 22.6) for which all three trees estimated very high mutation rates. All three trees generated mutation rate estimates for SL307; these overlapped one another (Verticall alignment: 4.2 substitutions/year [95% CI: 2.4 – 6.1]; Verticall distance: 4.4 substitutions/year [95% CI: 3.5 – 5.5]; Gubbins: 4.6 substitutions/year [95% CI: 3.2 – 6.0]; see **Tables S9–S11**) but were lower than published mutation rates estimated using BEAST for ST307, the dominant sequence type (ST) in this clone (6.6 substitutions/year [95% CI: 4.5 – 8.8]) [42]. Mutation rates estimated here for SL258 (which includes ST258, ST11, ST512, ST340, ST395 and >35 other STs) also overlapped (Verticall distance: 3.6 substitutions/year [95% CI: 3.2 – 4.0]; Gubbins: 3.2 substitutions/year [95% CI: 2.9 – 3.5]), but were lower than published estimates for ST258 (5.8 substitutions/year [95% CI: 4.6 – 7.0]) [41]. For other common multidrug resistant clones SL17, SL147, and SL15/14, only the Verticall distance method recovered temporal signal to allow divergence date estimation. Similar to SL307 and SL258, our mutation rate estimate for SL147 (4.0 substitutions/year [95% CI: 3.4 – 4.6]) was lower than the 6.2 substitutions per year published for the dominant clonal group (CG) in this sublineage, CG147 [44]. Conversely, the mutation rate estimate for SL17 (3.6 substitutions/year [95% CI: 2.9 – 4.2]) was higher than that previously estimated for ST17 (2.2 substitutions/year [95% CI: 1.3 – 3.2]) [43].

## Conclusions

While we consider Gubbins the gold-standard for detection of recombination within clonal lineages, for large datasets of thousands of genomes (where Gubbins fails to complete in a reasonable time frame), or analyses that extend beyond a single lineage such as species-wide or genus-wide variation, Verticall provides a useful solution to more efficiently and reliably identify recombination. Between the two approaches implemented in Verticall, the distance-based workflow (Verticall distance) showed superior performance, in terms of equivalence of results to Gubbins (Figures 3, 5 and 6, Table 1) or recovery of temporal signal (Figures 7, 8, S5, Table S7), compared with Verticall alignment and is therefore recommended in most cases. However, for large datasets, the pairwise distance-tree approach becomes inefficient due to the large number of comparisons required, and Verticall’s alignment-based approach is needed. Results with Verticall alignment are generally not as reliable as Verticall distance, but it did produce similar R^2^ and tree structures to Gubbins on the large *S.* Typhi dataset (Figures 4 and S6), and overlapping rate and date estimates for most *K. pneumoniae* clonal lineages tested (Figure 8), showing it is a reasonable solution.

## Author Contributions

**Conceptualization:** Ryan R Wick, Kathryn E Holt

**Data Curation:** Ryan R Wick, Erkison Ewomazino Odih

**Formal Analysis:** Ryan R Wick, Erkison Ewomazino Odih

**Methodology:** Ryan R Wick

**Software:** Ryan R Wick, Erkison Ewomazino Odih

**Writing – Original Draft Preparation:** Ryan R Wick, Erkison Ewomazino Odih

**Writing – Review & Editing:** Kathryn E Holt.

## Ethics Statement

Ethical approval was not required as only publicly available whole genome sequence data with no patient-level information were used in this study.

