## Supplementary Figures for "Verticall: A fast and robust tool for recombination detection in large-scale bacterial genomic datasets"

Figure S1: Comparison of the median and interpolated median pairwise distance estimates

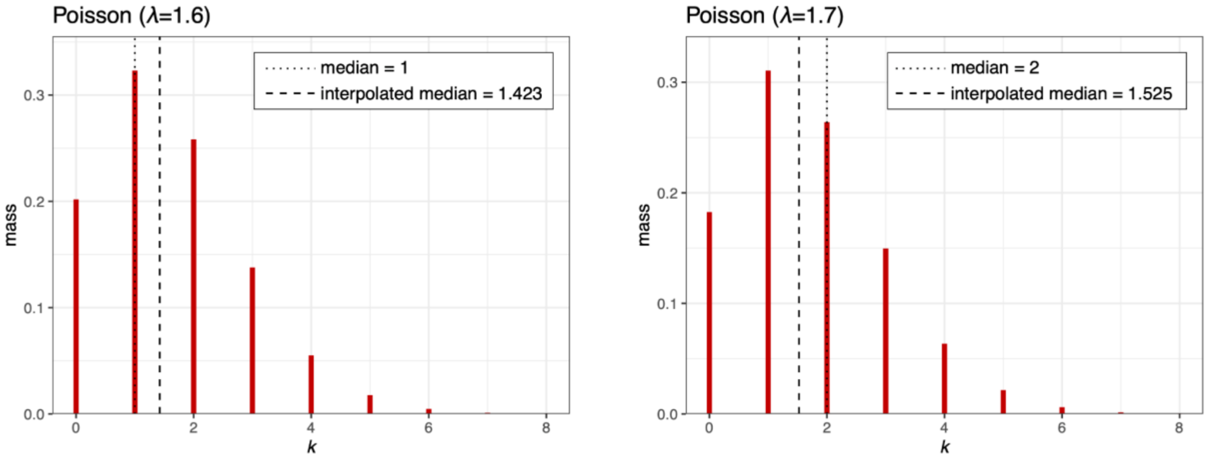

Figure S2: Comparison of Gubbins and Verticall distance trees for the *S. pneumoniae* PMEN1 lineage

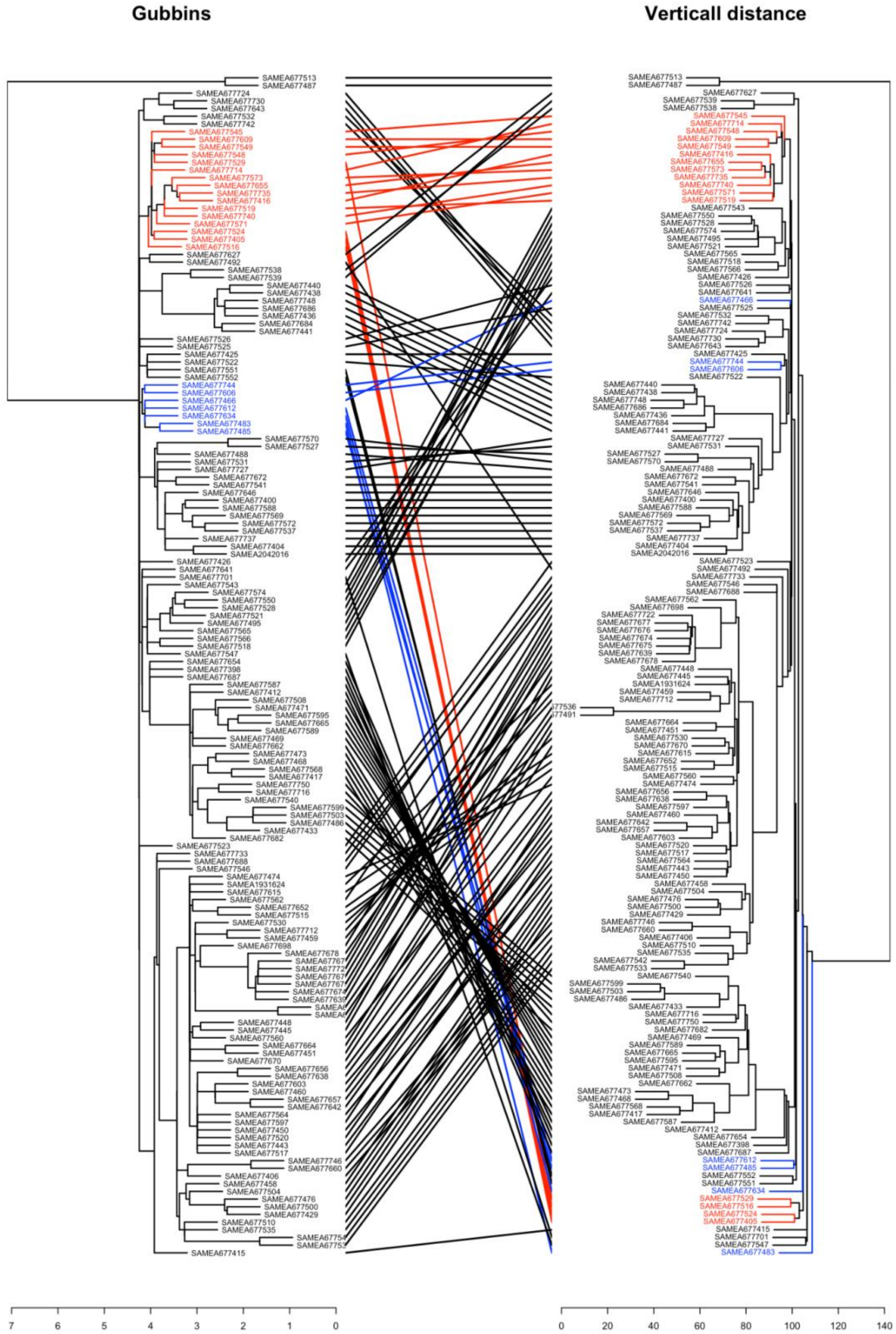

Figure S3: Comparison of ClonalFrameML and Verticall distance trees for the *E. coli* species-wide dataset

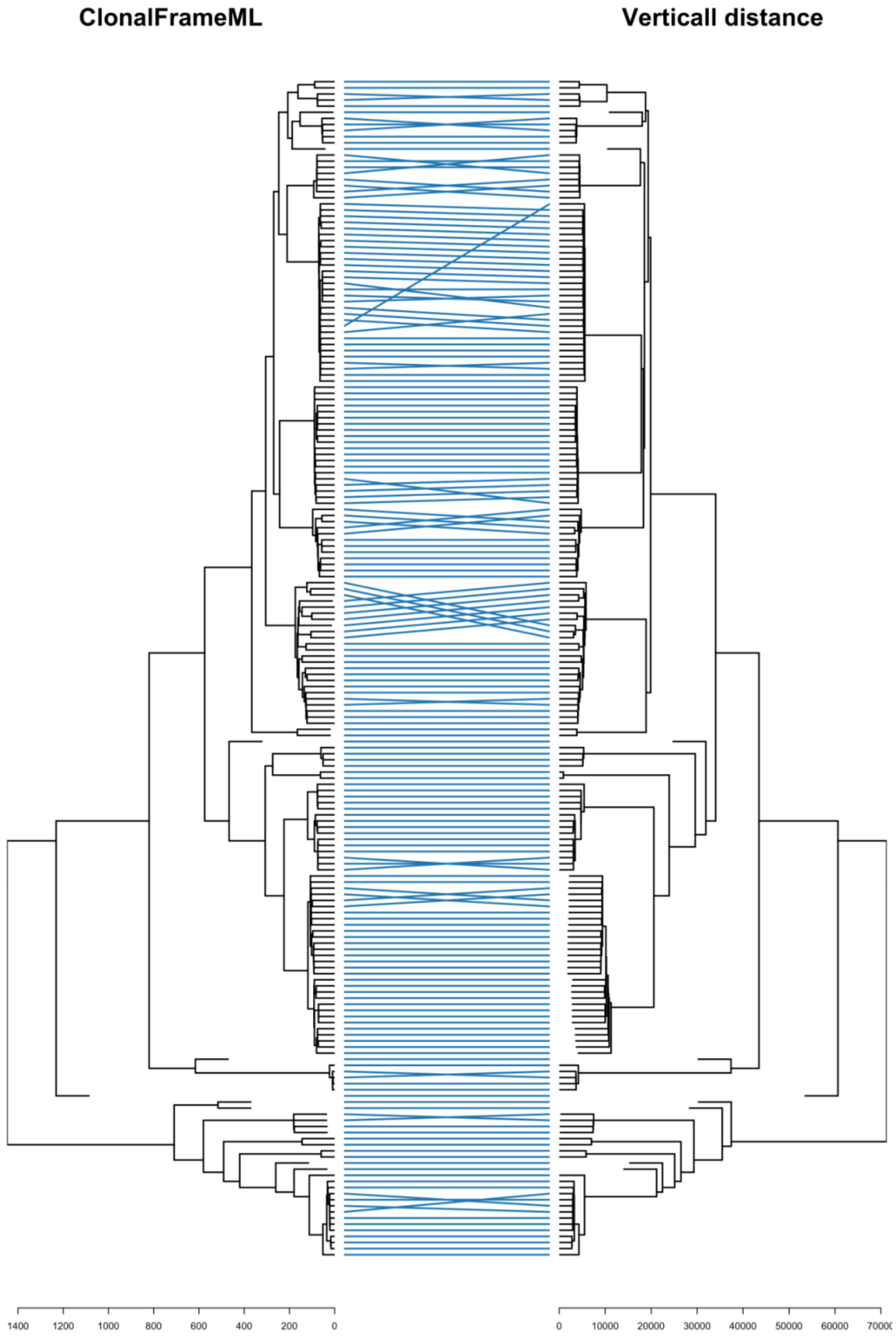

### Figure S4: Recombination filtering and phylogeny inference for the *Klebsiella* genus-wide sequences

Plots show the recombination blocks (right subpanels) and final trees (left subpanels) produced using the ClonalFrameML **(A)** and Verticall distance **(B)** workflows. Unfiltered trees generated using alignment-based **(C)** and distance-based **(D)** approaches are also shown. The tree branches are shaded to indicate the different *Klebsiella* species, as per the figure legend. Tree scales indicate number of substitutions. Each row in the recombination plot represents a single genome, and the columns correspond to nucleotide positions in a reference genome (*K. pneumoniae* 30660/NJST258_1 chromosome; GCF_000598005.1). In panel (A), the recombination blocks are coloured to indicate recombination regions predicted in internal (shared by multiple genomes) or terminal branches, as per the panel legend. In (B), the blocks shown are pairwise recombination blocks predicted by the Verticall distance workflow relative to other genomes in the dataset. Overlapping recombination blocks were merged and the blocks are coloured to indicate the number of genome pairs relative to which the recombination regions were detected. Reference-based positions for each block were determined by aligning each genome to the reference; blocks outside reference-aligned positions are not shown (n=18,353 / 40,538). **(B)** Alignment-based tree produced with no recombination filtering **(C)** Distance-based tree produced with no recombination filtering. The tree branches are shaded to indicate distinct *Klebsiella* species, as per the figure legend.

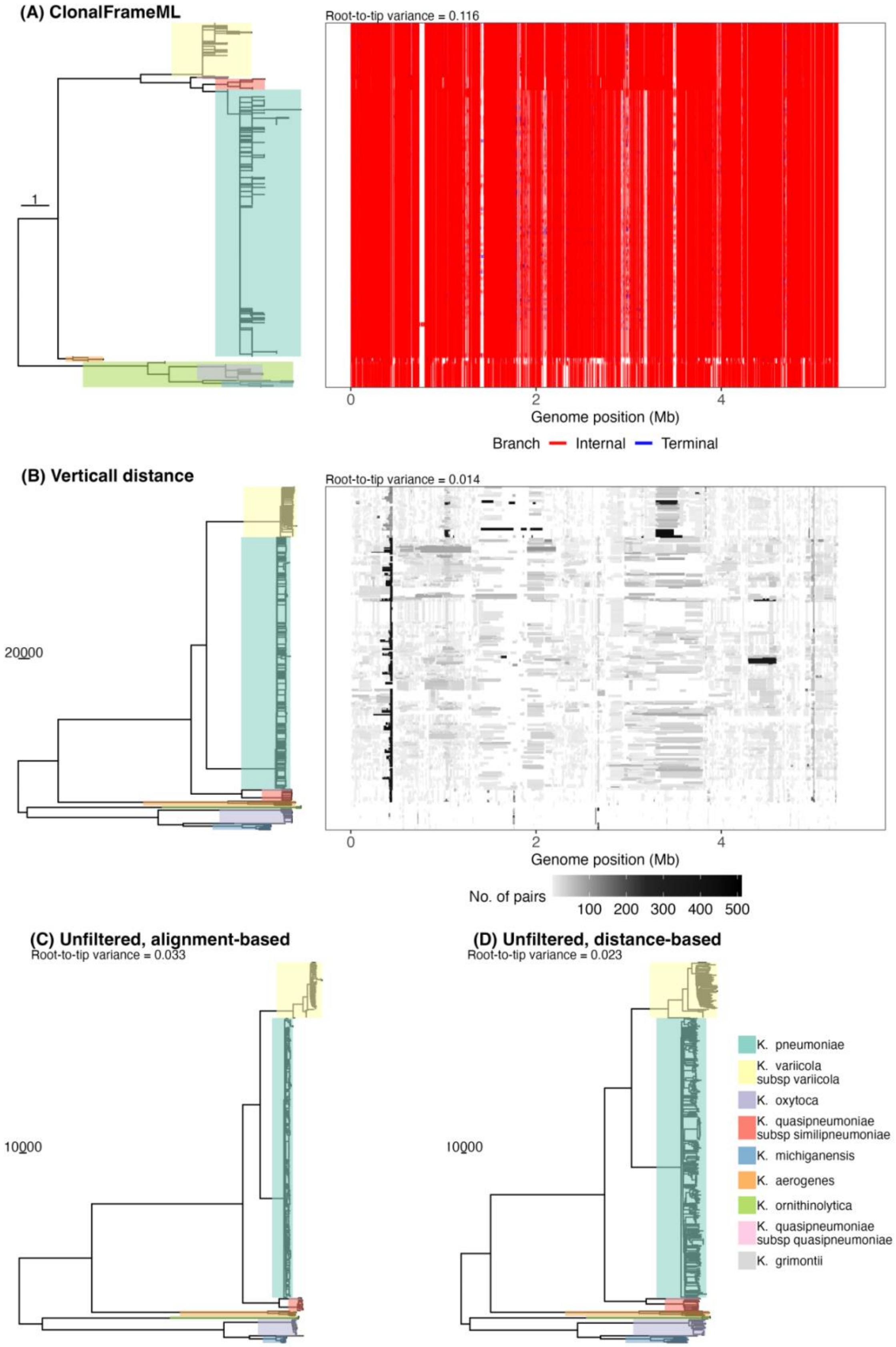

(A) ClonalFrameML
Root-to-tip variance = 0.116
1
0
2
4
Genome position (Mb)
Branch
Internal
Terminal
(B) Verticall distance
Root-to-tip variance = 0.014
20000
0
2
4
Genome position (Mb)
No. of pairs
100
200
300
400
500
(C) Unfiltered, alignment-based
Root-to-tip variance = 0.033
10000
(D) Unfiltered, distance-based
Root-to-tip variance = 0.023
10000
K. pneumoniae
K. variicola subsp variicola
K. oxytoca
K. quasipneumoniae subsp similipneumoniae
K. michiganensis
K. aerogenes
K. ornithinolytica
K. quasipneumoniae subsp quasipneumoniae
K. grimontii

Figure S5: Linear correlation between root-to-tip distances and sampling dates of *S. pneumoniae* PMEN1 lineage

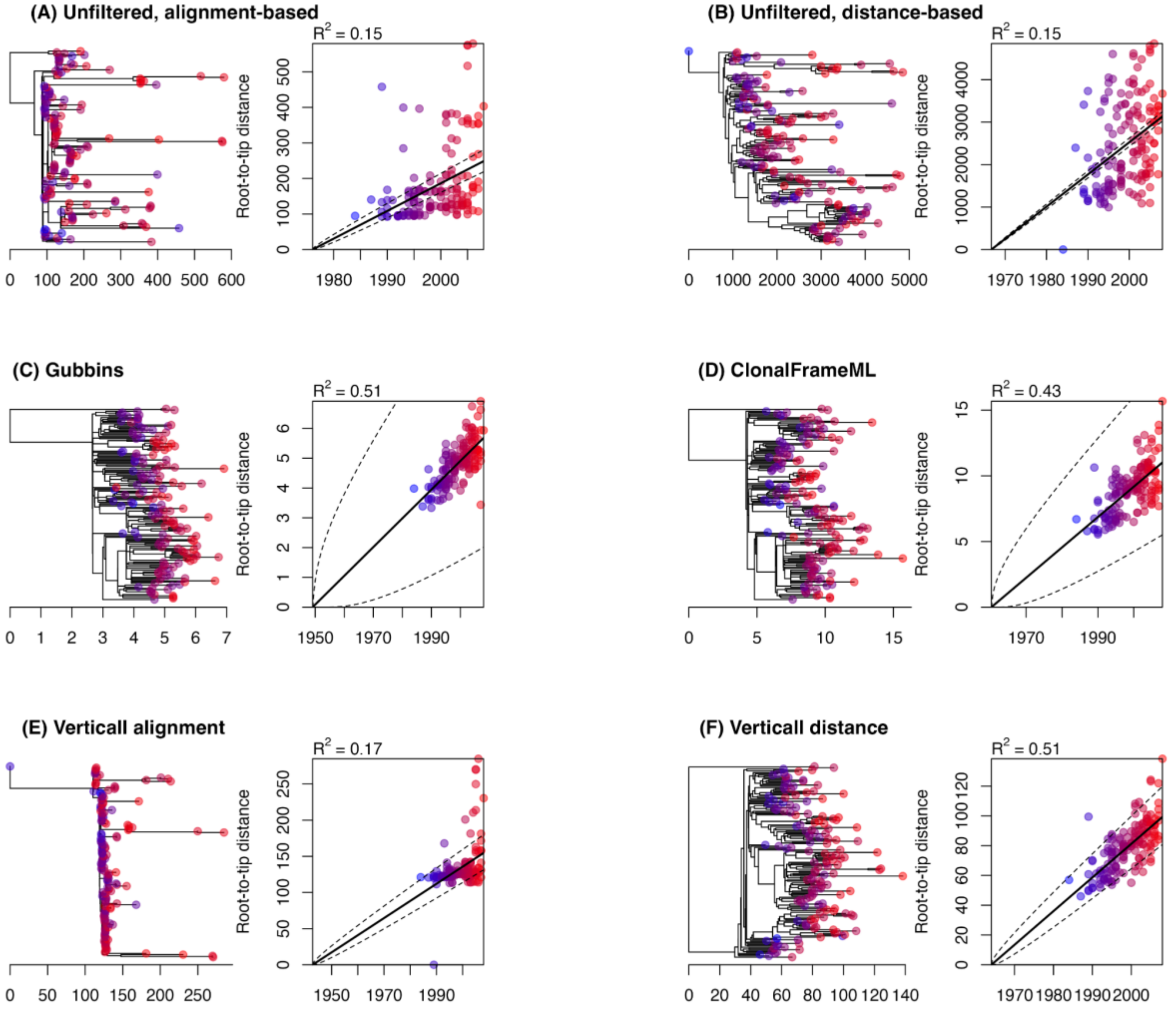

Figure S6: Linear correlation between root-to-tip distances and sampling dates of *S. enterica* Typhi H58 lineage

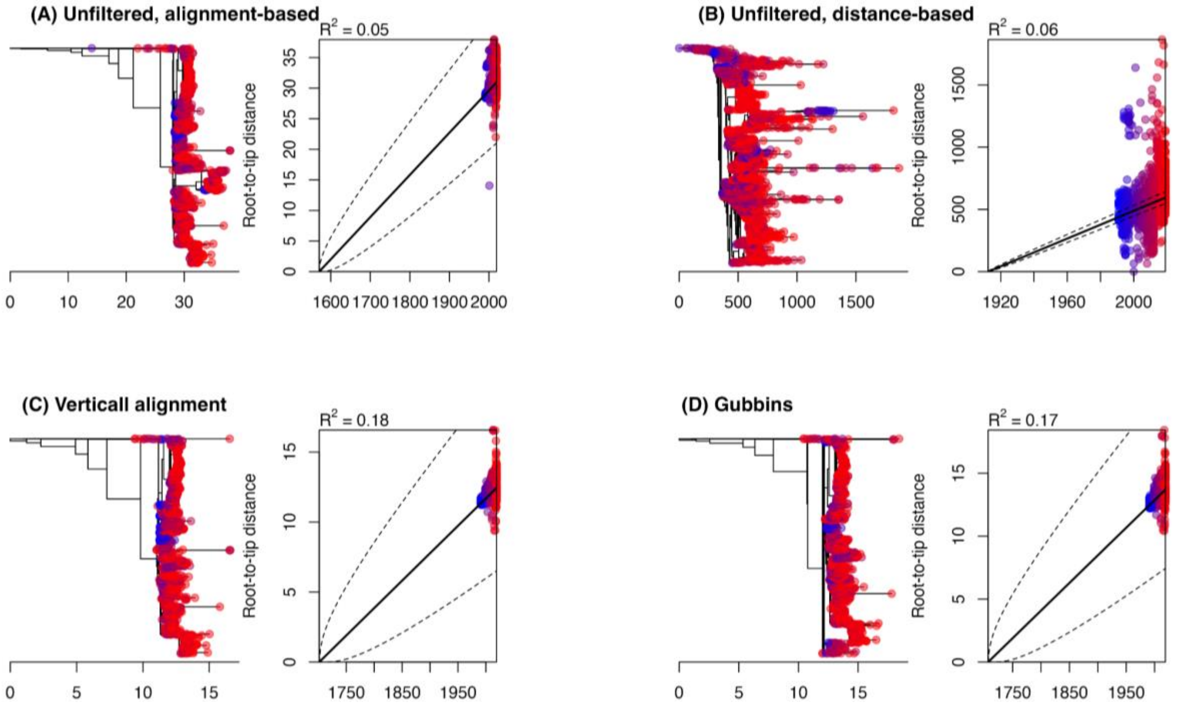

Figure S7: Convergence of divergence time estimation runs across 83 *Klebsiella pneumoniae* sublineage datasets

Plots show comparisons of convergence frequency using all three workflows and stratified by nucleotide divergence (A) and the number of genomes (B) in the datasets.

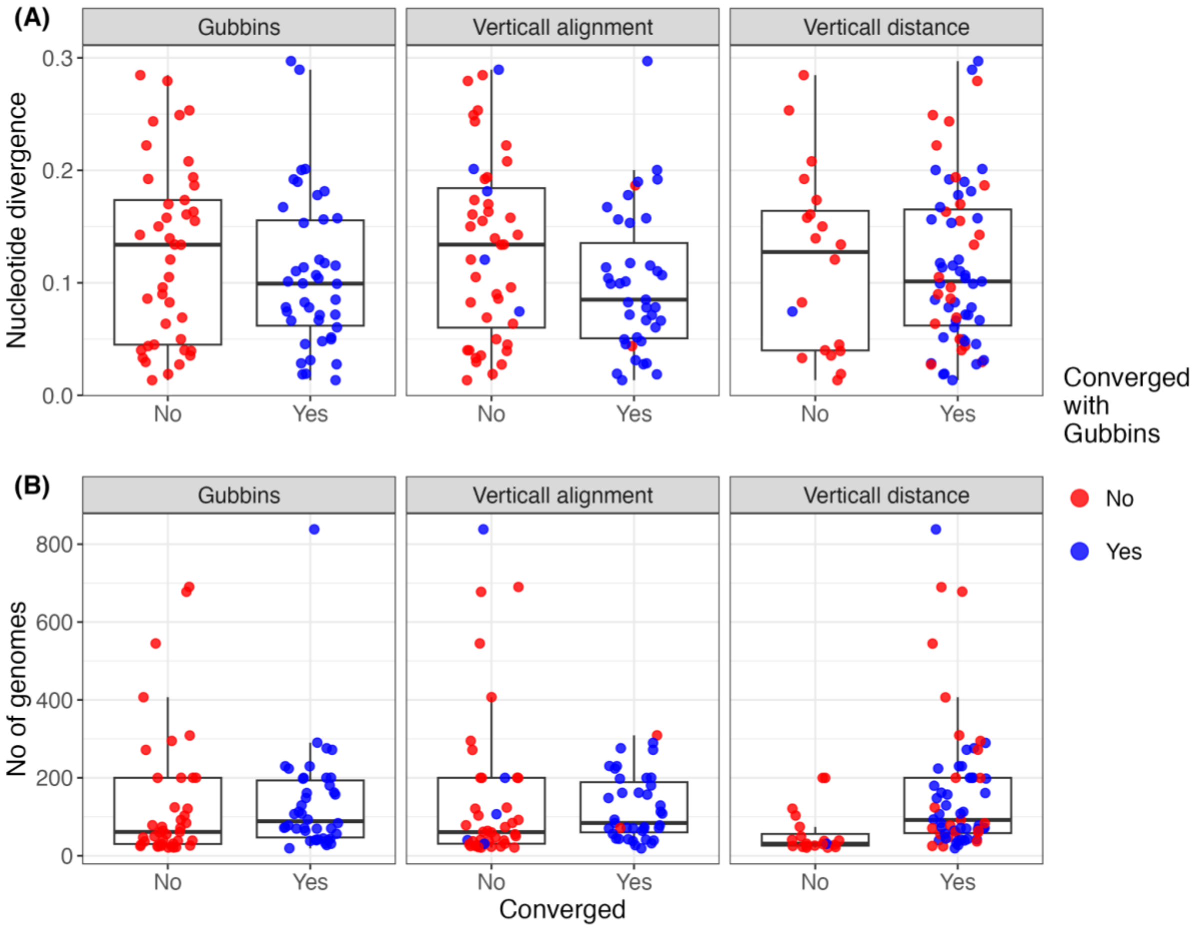
